# zingeR: unlocking RNA-seq tools for zero-inflation and single cell applications

**DOI:** 10.1101/157982

**Authors:** Koen Van den Berge, Charlotte Soneson, Michael I. Love, Mark D. Robinson, Lieven Clement

## Abstract

Dropout in single cell RNA-seq (scRNA-seq) applications causes many transcripts to go undetected. It induces excess zero counts, which leads to power issues in differential expression (DE) analysis and has triggered the development of bespoke scRNA-seq DE tools that cope with zero-inflation. Recent evaluations, however, have shown that dedicated scRNA-seq tools provide no advantage compared to traditional bulk RNA-seq tools. We introduce zingeR, a zero-inflated negative binomial model that identifies excess zero counts and generates observation weights to unlock bulk RNA-seq pipelines for zero-inflation, boosting performance in scRNA-seq differential expression analysis.

## Introduction

Transcriptomics has become one of the standard tools in modern biology to unravel the molecular basis of biological processes and diseases. One of the most common applications of transcriptome profiling is the discovery of differentially expressed (DE) genes, which exhibit changes in average expression levels across conditions.^1–3^ Over the last decade, RNA-seq has become the standard technology for transcriptome profiling enabling researchers to study average gene expression over bulks of cells.^4,5^ The advent of single cell RNA-seq (scRNA-seq) enabled high-throughput transcriptome profiling of single cells and disrupted research on developmental trajectories, cell-to-cell heterogeneity and the discovery of novel cell types, amongst others.^6–11^

In scRNA-seq, individual cells are first captured and the RNA is converted to cDNA in a reverse transcription step upon which vast amplification of the minute amount of starting material occurs prior to sequencing.^12^ Many sc RNA-seq protocols have been published to conduct these core steps,^13–18^ but despite these advances, sc RNA-seq data remains inherently noisy. Dropout events cause many transcripts to go undetected due to inefficient cDNA polymerisation, amplification bias or low sequencing depth, leading to excessive zero counts, as compared to bulk RNA-seq data.^18,19^ The presence of dropouts suggest two different types of zeros in scRNA-seq data: biological zeros, when a gene is simply not expressed in the cell, and excess zeros, when a gene is expressed in the cell but it was not observed for technical reasons other than a low sequencing depth. In the count data analysis literature this is also referred to as zero-inflation. In addition, scRNA-seq counts are inherently more variable than bulk RNA-seq data because the transcriptional signal is not averaged across thousands of individual cells (Supplementary Figure 1), making cell-to-cell heterogeneity, cell type mixtures and stochastic expression bursts important contributors to between-sample variability.^7,20^

**Figure 1:**
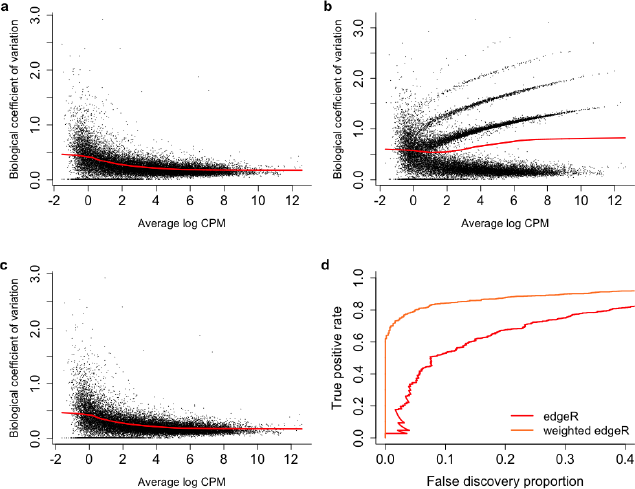
Zero-inflation results in overestimated dispersions and jeopardizes power to discover differential expression. The biological coefficient of variation (BCV) is the square root of the NB dispersion parameter. (a) The BCV plot of a regular bulk RNA-seq experiment. Dispersion estimates generally show a smooth decrease for genes with higher expression. (b) Simulating zero-inflation by randomly introducing 5% excess zero counts inflates dispersion estimates for the genes with excess zeros. This significantly distorts the estimated mean-variance relationship, as represented by the red line. (c) Downweighting excess zeros by assigning the introduced zeros a weight of zero in dispersion estimation recovers the original mean-variance trend. (d) False discovery proportion - true positive rate (FDP-TPR) performance curves on the zero-inflated data shows that the performance of edgeR (red curve) is deteriorated in a zero-inflated setting due to an overestimation of the dispersion parameter. However, assigning the introduced zeros a weight of zero in the dispersion estimation and model fitting results in a dramatic performance boost. Hence, it is the key to unlocking RNA-seq tools for zero-inflation.

In RNA-seq applications, abundances are typically estimated by using counts, which represent the number of sequencing reads mapping to an exon, transcript or gene. Popular RNA-seq DE tools like edgeR^2^ and DESeq2^1^ assume a negative binomial count distribution across biological replicates, while limma-voom^3^ uses linear models to model log-transformed counts and observation-level weights to account for the mean-variance relationship of the count data. These bulk RNA-seq tools have can also be applied for scRNA-seq DE analysis^21^. However, dropouts and high variability in scRNA-seq data raised concerns about the utility of existing bulk RNA-seq tools for scRNA-seq data analysis. This has triggered the development of novel dedicated tools, which typically introduce an additional component to account for excessive zero counts through, for example, zero-inflated (scde^22^) or hurdle models (MAST^19^). However, Jaakkola et al. (2016)^23^ and Soneson & Robinson (2017)^24^ have recently shown that these bespoke tools do not reveal systematic benefits over standard RNA-seq tools in scRNA-seq applications.

We argue that standard RNA-seq tools, however, still suffer in performance due to zero-inflation with respect to the negative binomial distribution. We illustrate this using biological coefficient of variation (BCV) plots,^25^ which visualize the mean-variance relationship of the counts. Note, that the BCV plots of scRNA-seq datasets contain striped patterns (Supplementary Figure 2 for scRNA-seq datasets subsampled to ten samples), that are indicative for genes with few positive counts (Supplementary Figure 3) and very high dispersion estimates. Randomly adding zeros to bulk RNA-seq data, likewise consisting of ten samples, also results in very similar striped patterns (Figure 1). The negative binomial models implemented in DESeq2 and edgeR will thus accommodate excess zeros by overestimating the dispersion parameter, which jeopardizes the power to discover differential expression in the presence of zero-inflation. However, a correct identification of the introduced excess zeros and downweighting them by assigning a weight of zero in dispersion estimation and model fitting reconstructs the original mean-variance relationship (Figure 1c), recovering the power to detect differential expression (Figure 1d). Hence, identifying and downweighting excess zeros provide the key to unlock RNA-seq tools for scRNA-seq differential expression analysis.

**Figure 2:**
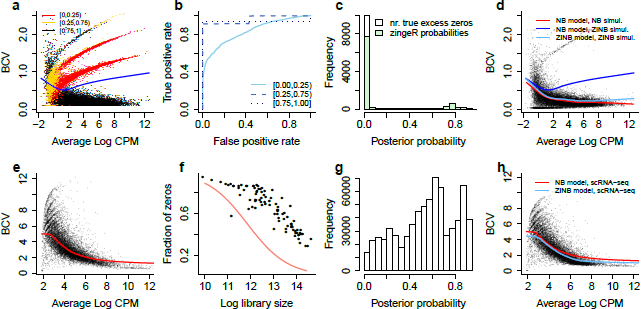
Zero-inflation distorts the mean-variance trend in (sc)RNA-seq data but is correctly identified by the ZINB model. The top panels represent simulated RNA-seq data while the bottom panels represent the scRNA-seq dataset from Islam et al. (2011).^16^ The biological coefficient of variation (BCV) is the square root of the NB dispersion parameter. (a) Simulated RNA-seq dataset for a two-group comparison with five samples in each group. Randomly replacing 5% of the expression counts with zeros induces zero-inflation and distorts the mean-variance trend by overestimating the dispersion parameter. The colors represent the average zingeR posterior probability for all zeros from a gene. (b) ROC curve for the correct identification of excess zeros, as stratified by the average log CPM. A very good classification precision is obtained for genes with moderate and high expression, while the identification is harder on genes with low expression, since these genes often have higher dispersion estimates. (c) Histogram of zingeR posterior probabilities for the excess zeros in the zero-inflated bulk RNA-seq dataset. The white bar at 0 represents the number of introduced excess zeros. Most excess zeros are identified as such by zingeR, see also Supplementary Figure 4. (d) Effectively downweighting excess zeros using the posterior probabilities recovers the original mean-variance trend and inference on the NB count component will now no longer be biased because of zero-inflation patterns, illustrating zingeR's ability to account for excess zeros. (e) BCV plot for the Islam dataset^16^ shows that higher variability is observed in scRNA-seq data as compared to bulk RNA-seq data. As in (b), zero-inflation induces striped patterns in the BCV plot leading to an overestimation of the dispersion parameter of the count component. (f) The sequencing depth of a cell is related to the abundance of zeros, information used by zingeR to identify excess zeros in scRNA-seq datasets when fitting the ZINB model. The pink curve represents the estimated marginal mean on excess zeros for a cell and the difference between the curve and the datapoints represents the estimated expected fraction of zeros that belong to the count component. (g) zingeR posterior probabilities for all zeros in the Islam dataset identify both NB and excess zeros. However, due to the increased noise in scRNA-seq datasets, some zeros are harder to classify as compared to bulk RNA-seq data. (h) Using the zingeR posterior probabilities as observation weights results in lower estimates of the dispersion parameter, unlocking powerful differential expression analysis with standard RNA-seq DE methods.

**Figure 3:**
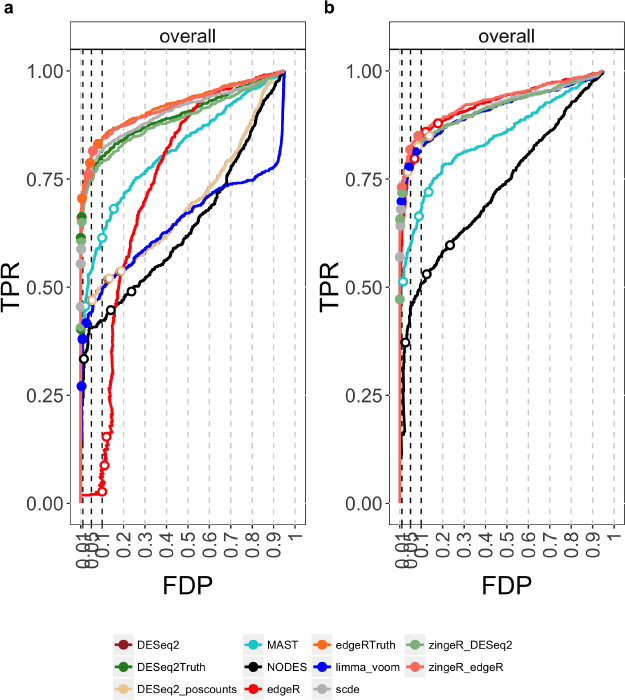
Comparison of methods on simulated RNA-seq data. The left panel (a) shows performance curves on the simulated dataset where 5% of the counts were set to zero. Conventional RNA-seq methods break down due to zero-inflation while most scRNA-seq methods perform reasonably, scde seems to have a good performance in a moderate zero-inflation setting, however it provides very conservative FDR control as suggested by its FDR working points. zingeR_edgeR outperforms all other methods and in fact performs close to an edgeR analysis based on the truth, where the introduced zeros are effectively downweighted by setting their weights to zero, showing that zingeR_edgeR correctly identified most excess zeros. A similar result is observed for zingeR_DESeq2. Right panel (b) shows the performance of the evaluated methods on simulated RNA-seq data, suggesting a superior performance of zingeR_edgeR and providing evidence that the method performs well in non zero-inflation settings. Note, that the working points for zingeR_DESeq2 are more conservative as compared to DESeq2 due to the use of the t-distribution instead of the Gaussian distribution for the Wald test.

We therefore propose `zingeR` (Zero Inflated Negative binomial Gene Expression in R), a tool for scRNA-seq DE analysis using a zero-inflated negative binomial (ZINB) distribution. `zingeR` efficiently identifies excess zeros and provides observation weights to unlock bulk RNA-seq pipelines for zero-inflation. `zingeR` is shown to outperform competing methods on simulated RNA-seq and simulated scRNA-seq datasets. We also demonstrate `zingeR’s` gain in performance in a case study on a differential expression analysis between neuronal cell types using publicly available data. The method and a novel simulation framework for scRNA-seq data are incorporated in the R package `zingeR` and are available at https://github.com/statOmics/zingeR.

## Results

### zingeR extends bulk RNA-seq tools towards zero-inflation

In this manuscript, we argue that standard RNA-seq tools applied in scRNA-seq applications still suffer from zero-inflation with respect to the negative binomial distribution. We propose zingeR, a tool for scRNA-seq DE analysis using a zero-inflated NB (ZINB) distribution, a two component mixture between a point mass at zero (δ) and a NB distribution (*f_NB_*)

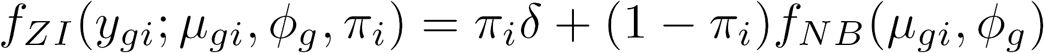

with *y_gi_* the expression counts for gene *g* in sample *i, π_i_* the mixture probability for an excess zero count, *μ_gi_* and *ϕ_g_* respectively the negative binomial mean and dispersion parameters. zingeR uses the EM-algorithm for fitting the mixture distribution, where we use the association between the cell’s sequencing depth and zero abundance to estimate *π_i_*. Upon convergence, zingeR produces posterior probabilities that a zero belongs to the count component

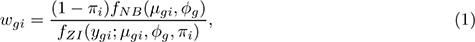

which can be used as observation weights in the analysis. We build upon edgeR to fit the ZINB model and assess DE using the negative binomial component of the mixture.

As in the introduction, we will demonstrate the problem and the solution provided by the zingeR method using BCV plots. We have already noted that adding zeros to bulk RNA-seq data results in striped patterns (Figure 1b, Figure 2a), that are indicative for genes with few positive counts (Supplementary Figure 3) and very high dispersion estimates. The zingeR model, however, identifies many introduced excess zeros as such (Figure 2a-c), by classifying them to the zero-inflation component of the ZINB mixture distribution. As expected, distinguishing biological from excess zeros is harder for genes with a lower expression, since those genes often have higher dispersion estimates (Figure 2b). Using zingeR posterior probabilities as observation-level weights in edgeR recovers the original BCV plot and mean-variance trend (Figure 2d), illustrating zingeR’s ability to account for zero-inflation. Hence, they provide the key to unlocking standard RNA-seq tools for zero-inflation. The BCV plot of the scRNA-seq Islam dataset (Figure 2e) shows a similar striped pattern as in the zero-inflated RNA-seq data. For scRNA-seq data, however, we also observed a trend between zero abundance and the cell’s sequencing depth (Supplementary Figure 4), information that we integrate in the zingeR model component for excess zeros (Figure 2f). Moreover, the data are more variable and the zero-inflation pattern is more subtle than in the RNA-seq example, resulting in a higher classification uncertainty of the zeros (Figure 2g). However, also in scRNA-seq applications, a decrease in dispersion estimates is achieved (Figure 2h), suggesting that zero-inflation patterns were indeed present and have been accounted for.

**Figure 4:**
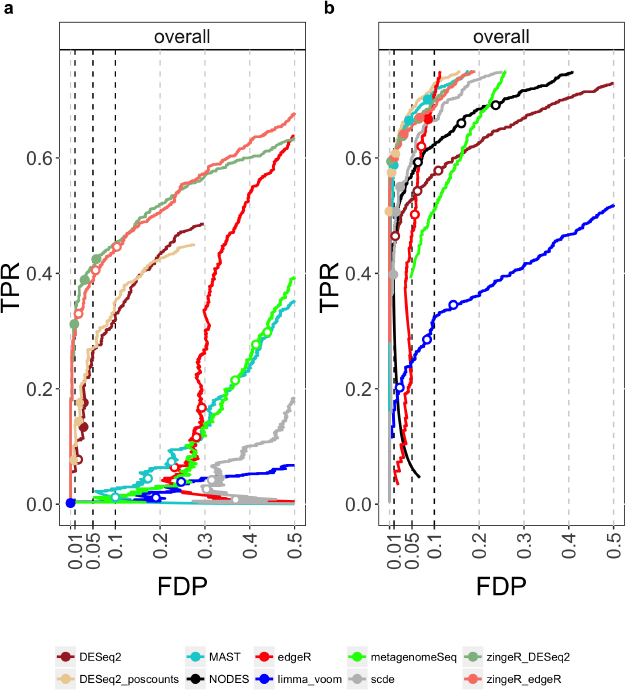
Comparison of methods on simulated scRNA-seq data. (a) Performance on scRNA-seq data based on Islam simulation. (b) Performance based on Trapnell simulation. The zingeR workflows clearly outperform other methods in case of severe zero-inflation (a) and are among the best performers in the Trapnell simulation with few excess zeros (b). Note, that the positive counts normalization provides an enormous boost in performance for DESeq2 in the Trapnell simulation. Dedicated methods scde and metagenomeSeq specifically developed to deal with excess zeros are dominated in both simulations by zingeR workflows and by DESeq2 with positive counts normalization. DESeq2 curves in panel (a) are cut-off due to NA p-values as a result of independent filtering. Full FDP-TPR curves are provided in Supplementary Figure 9.

### RNA-seq simulation study

First, we evaluate zingeR in an RNA-seq context. We use zingeR weights in conjunction with edgeR (zingeRedgeR) and DESeq2 (zingeR_DESeq2) and compare them to state-of-the-art bulk RNA-seq tools edgeR,^2,25^ DESeq2^1^ with default and positive counts normalization (see Online Methods) and limma-voom;^3^ scRNA-seq dedicated tools scde,^22^ MAST^19^ and NODES;^26^ and metagenomeSeq^27^ developed for zero-inflation in metagenomics applications. We use the framework from Zhou et al. (2014)^28^ to estimate gene-wise parameters from the Bottomly dataset^29^ and simulate RNA-seq counts for a two-group comparison according to a gene-wise negative binomial distribution where the means and dispersions are jointly sampled to respect the original mean-variance relationship. In a first scenario, we evaluate the methods in a zero-inflated RNA-seq setting, by randomly replacing 5% of all counts with excess zeros. The false discovery proportion - true positive rate (FDP-TPR) curves (Figure 3) clearly illustrate that all conventional RNA-seq DE tools break down since the excess zeros inflate dispersion estimates (Figure 2b) and thus compromise performance. zingeR, however, correctly identifies excess zeros, and thus recovers the original mean-variance relationship, boosting performance for the differential expression analysis. zingeR_edgeR has superior performances in this simulation, but is closely followed by zingeR_DESeq2 and scde. However, scde provides very conservative FDR control as suggested by its FDR working points. All three methods use a zero-inflated negative binomial distribution to model the counts, and convincingly outperform the other tools. MAST also outperforms the bulk RNA-seq tools, but is inferior to the methods using zero-inflated distributions. Furthermore, zingeR_edgeR and zingeR_DESeq2 even attain a similar performance as an edgeR or DESeq2 analysis using the ground truth, i.e. by assigning excess zeros a zero weight, clearly demonstrating the correct identification of excess zeros by the zingeR algorithm. Moreover, in the absence of zero-inflation, the performance of zingeR_edgeR and zingeR_DESeq2 is not deteriorated (Figure 3) and they converge to a regular edgeR (DESeq2) analysis. Hence, adopting the zingeR methods will not harm the analysis in any case.

### scRNA-seq simulation study

The RNA-seq simulation study has shown that tools adopting zero-inflated distributions have superior performances in an RNA-seq setting with excess zeros. scRNA-seq data, however, are noisier than RNA-seq data and the excess zeros do not occur completely at random. Therefore, we provide a scRNA-seq data simulation paradigm that retains gene-specific characteristics as well as global associations across all genes. More specifically, we estimate the dataset-specific associations between zero abundance with the sequencing depth and average expression rates and explicitly model this in our simulation framework (Supplementary Figures 4-5). The scRNA-seq simulation is based on two datasets: the Islam^16^ mouse dataset, which compares 48 embryonic stem cells to 44 embryonic fibroblasts in mouse, and the 48h and 72h timepoints of the human Trapnell^30^ dataset, comparing differentiating human myoblasts at the 48h (85 cells) and 72h (64 cells) timepoints. The datasets differ in their extent of zero-inflation (Supplementary Figure 6) and provide a basis for method evaluation and comparison at different degrees of zero-inflation that occur in practice. The simulated datasets successfully mimic the characteristics of the original datasets (Supplementary Figures 7, 8), suggesting good quality of the simulated data. Figure 4 (Supplementary Figure 9) illustrates that many methods break down on the simulated Islam dataset due to a high degree of zero-inflation. Surprisingly, even methods specifically developed to deal with excess zeros like MAST, scde and metagenomeSeq suffer from poor performances. The DESeq2 methods, however, are able to cope with a high degree of zero-inflation. In general, it is a good strategy to disable the imputation step in DESeq2, since it deteriorates its performance in scRNA-seq data (Supplementary Figure 10). The zingeR models dominate all competitors in terms of sensitivity and specificity, providing high power and good FDR control. These results persist even when simulating DE with high fold changes (> 3) (Supplementary Figure 11). Although scde had good performances in the bulk RNA-seq simulations it has low power and bad FDR control in a high zero-inflation setting. Note, however, that the remaining methods also suffer from poor FDR control.

**Figure 5:**
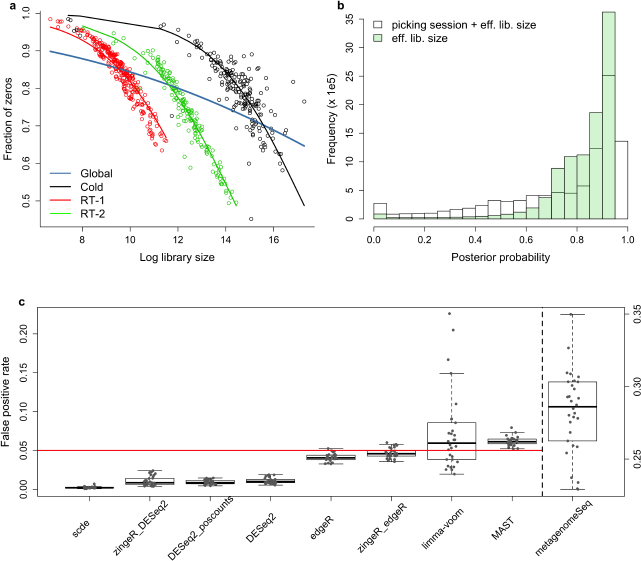
Case study on neuronal cells. (a) The association of zero abundance with sequencing depth. The three different picking sessions differ in their sequencing depth, causing an attenuated global relationship (blue line). However, accounting for the batch effect in zingeR’s zero-excess model, allows for a correct model of sequencing depth with zero abundance. (b) The distribution of posterior probabilities with (white) and without (green) including the batch effect in the zero-excess model. Including the batch effect results in many more zeros being identified as truly zero excess (i.e. the higher bar near a posterior probability of zero) and negative binomial zeros, hence increasing classification certainty. (c) False positive rate (FPR) evaluation on 30 mock comparisons across all cell types. Note, that a different scale is used for metagenomeSeq.

Since zero-inflation is fairly modest in the Trapnell dataset, most methods perform better than in the Islam simulation (Figure 4). zingeR_edgeR, zingeR_DESeq2, MAST and DESeq2 with positive counts normalization, outperform the remaining methods in this simulation in terms of sensitivity and provide good FDR control. scde is their closest competitor, however the method is again overly conservative. Notably, DESeq2 has very liberal FDR control, but the positive counts normalization results in a dramatic performance boost. The scRNA-seq method NODES provides good sensitivity, but it also suffers from a very liberal FDR control. Note, that the performance of all methods is still lower than in the bulk RNA-seq simulation due to the high level of noise associated with scRNA-seq experiments.

### Case study: differential expression between neuronal cell types

Finally we apply zingeR to a publicly available scRNA-seq dataset for neuronal cell types in mouse.^11^ In this experiment, cells from the dorsal root ganglion were robotically picked and the 5’ end of the transcripts were sequenced. After quality control, the authors considered 622 cells which were classified in eleven neuronal cell type categories. The authors acknowledge the existence of a batch effect that is related to the picking session of the cells, where all cells were picked in three separate picking sessions. We find that the batch effect is not only associated with expression but also influences the association of sequencing depth with zero abundance (Figure 5a).^31^ Large differences in sequencing depth between batches (picking sessions) causes an attenuation of the global association across all batches (Figure 5a). We therefore add the batch effect as a covariate in both the zingeR count model and zero-excess model. Upon correction for batch effects, a better identification of excess zeros is achieved (Figure 5b) with a higher classification certainty. This shows the generality of the zingeR method: both the count component as well as the zero-excess model component can be modelled in a very flexible way, providing an optimal assessment of differential expression while accounting for any factor that can improve the identification of excess zeros. The authors identify genes that characterize each cell type by comparing the expression for each neuronal cell type with the average expression of the remaining cell types. In the original manuscript, the analysis was performed with scde^22^ with a batch correction procedure that accounts for the picking sessions. Supplementary Table 1 shows that all methods provide higher numbers of significant genes than scde, and zingeR_edgeR has the highest number of significant genes across all methods. This is in agreement with the simulation studies, where zingeR_edgeR typically provides highest sensitivity. However, since a higher number of differentially expressed genes may arise due to a higher number of false positives, we evaluated the false positive rate in this dataset using 30 random 45 vs. 45 mock comparisons. In every condition, 15 cells from each of the three picking sessions are randomly selected over all cell types, thereby controlling for potential confounding by this batch variable, and we test for significance of the mock variable. The FPR is controlled by both zingeR variants, suggesting that the high number of significant genes for the zingeR models is not due to a higher fraction of false positives in the significance list (Figure 5c). The mock comparison also shows that limma-voom is too liberal in some evaluations and MAST consistently provides somewhat liberal results, while especially metagenomeSeq is extremely liberal. In addition, a uniform p-value distribution is observed for zingeR_edgeR in the mock comparison, while there seems to be issues with the null distribution of the test statistics from DESeq2 methods and scde, which produce too conservative p-values (Supplementary Figure 12).

## Discussion

We used default bulk RNA-seq normalization procedures and adopted positive counts normalization for DESeq2,^32^ which has been shown to boost its performance. Novel normalization procedures have been developed for scRNA-seq data analysis (e.g. scran^33^), but a thorough comparison of normalization methods falls outside the scope of this contribution. The zingeR implementation, however, allows the user to supply custom normalization factors, which opens the zingeR data analysis workflow towards any normalization method that produces normalization factors or offsets.

In all simulations, we have used the zingeR count component for inference, using either edgeR or DESeq2. However, zingeR’s posterior probabilities can also be used to unlock other standard RNA-seq workflows in zero-inflation situations. Supplementary Figure 13 shows that zingeR observation weights also boost performance of limma-voom in an scRNA-seq context, where we combine heteroscedastic weights with the posterior probabilities. Similar to the default limma-voom method, the zingeRlimma-voom implementation suffers from a liberal FDR control.

Our simulations complement the findings of Jaakkola et al. (2016)^23^ and Soneson & Robinson (2017),^24^ but additionally suggest that the performance of dedicated scRNA-seq methods depends on the degree of zero-inflation. Although MAST, metagenomeSeq and scde were explicitly developed to address excess zeros, they suffer from poor performance in a high zero-inflation setting, as is demonstrated in the Islam simulation study.

The method was demonstrated on scRNA-seq protocols relying on standard read counting. Recently, unique molecular identifiers (UMI) have been proposed to reduce the measurement variability across samples.^15^ In UMI-based protocols, transcripts are labeled with a small random UMI barcode prior to amplification. After amplification and sequencing, one then counts the number of unique UMIs found for every transcript, which then corresponds to the number of sequenced UMI-labeled transcripts. It has previously been shown^34^ that UMI-tagged data follow a negative binomial distribution. Hence, the zingeR methods will also provide good results for UMI-based data as they have the desirable property to converge towards a regular edgeR (DESeq2) analysis in the absence of zero-inflation. The latter is an important property and demonstrates zingeR’s broad applicability.

## Conclusions

In summary, we provide a realistic simulation framework for single cell RNA-seq data and introduce a novel tool zingeR that successfully identifies excess zeros related to dropout events in scRNA-seq experiments. We confirmed that state of the art scRNA-seq tools do not improve upon common RNA-seq tools for differential expression analysis of single cell RNA-seq experiments. The zingeR workflows, however, outperform current methods and have the merit to converge to conventional RNA-seq analyses in the absence of zero-inflation. Standard inference is provided by the count component of the ZINB model and our tool produces posterior probabilities that can be used as observation-level weights by conventional RNA-seq tools. Hence, zingeR has the promise to unlock traditional RNA-seq DE workflows for zero-inflated data and will assist researchers, data analysts and developers to improve the power to detect DE in the presence of excess zeros.

## Methods

### Negative binomial model

Let *Y_gi_* be the read counts for gene *g* in sample *i*. Many RNA-seq differential expression (DE) analysis tools^1,25,35^ assume the read counts to follow a negative binomial (NB) distribution

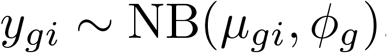

with *μ_gi_* the expected count in sample *i* and *ϕ_g_* the NB dispersion parameter. We adopt the negative binomial parametrization *Y* ~ NB(*μ, ϕ*), then *E*(*Y*) = *μ* and Var(*Y*) = *μ* + *ϕμ*^2^. Due to the low sample size in common RNA-seq experiments dispersion estimates as estimated with standard maximum likelihood theory are often unreliable and empirical Bayes methods^36,37^ are used to borrow information across genes. The negative binomial distribution is then embedded in a generalized linear model (GLM) with a log-link to model the mean *μ_gi_* of experiments with complex designs, i.e.

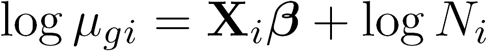

with **X**_*i*_ the covariates for observation *i, β* the model parameters of the linear predictor, and log *N_i_* an offset used for normalization, e.g., to correct for differences in sequencing depth and composition.^38^

### The zero-inflated negative binomial model (ZINB)

The major difference between scRNA-seq and bulk RNA-seq experiments is arguably the high abundance of zeros in scRNA-seq datasets. Traditionally, excess zeros are dealt with by the use of hurdle models or zero-inflated distributions, as recently proposed by Finak et al. (2015),^19^ Kharchenko et al. (2014)^22^ and Paulson et al. (2013).^27^ A zero-inflated count distribution *f_ZI_* is a two component mixture distribution between a point mass at zero δ and a count distribution *f_count_*, in our case the negative binomial distribution *f_NB_*:

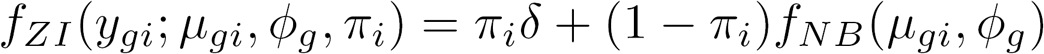

with *π_i_* the mixture parameter indicating the probability for a count to be an excess zero in sample *i*. The model parameters {*μ_gi_, ϕ_g_, π_i_*} can be estimated with maximum likelihood. However, no closed form solutions exist and we develop an expectation maximization (EM) algorithm for high throughput data. Note, that the ZINB model provides posterior probabilities that a count *y_gi_* belongs to the count component given the observed **X**_*i*_ and *N_i_*, which play a central role in our EM algorithm and can be used as observation weights in regular RNA-seq workflows. Let *Z_gi_* be an indicator variable, where *z_gi_* = 1 if *y_gi_* belongs the zero-inflation component and *z_gi_* = 0 if *y_gi_* originates from the count component, then the posterior probabilities are given by

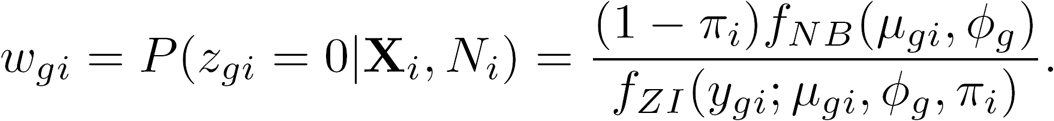

### EM algorithm

The EM-algorithm recasts the mixture model into a missing data problem by introducing the latent variable *Z_gi_*, which is assumed to follow a Bernoulli distribution: *Z_gi_* ~ *B*(*π_i_*). Hence, the likelihood

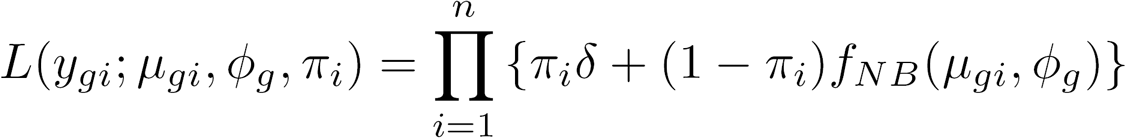

can be augmented with the *z_gi_* resulting in the joint likelihood

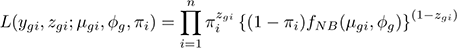

which in turn allows an efficient factorization by conditioning on *z_gi_*. The EM-algorithm iterates over an expectation (E) and maximization (M) step until convergence. In the E-step the expected joint log likelihood is calculated given the data and the current values of the parameter estimates, i.e. the parameter estimates in the previous iteration 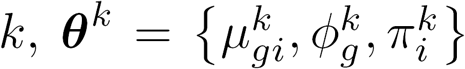. In the M-step the expected log-likelihood is maximized to update the parameter estimates. For our mixture model, the expected joint log-likelihood *l*(*y_gi_, z_gi_*) given the data and the current parameter estimates equals

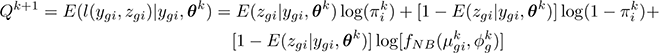

which shows that calculating *Q*^*k*+1^ only involves replacing *z_gi_* by its conditional expectation 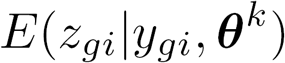 after which *Q*^*k*+1^ can be maximized over the mixture distribution parameters. In this expression, 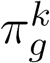 represents the parameter estimate for gene *g* in step *k*, and similar notation is used for the other parameters.

1. **E-step:** Calculate *Q*^*k*+1^ using the conditional expectations 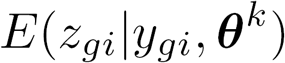, which are the posterior probabilities for counts to belong to the zero-inflation component:

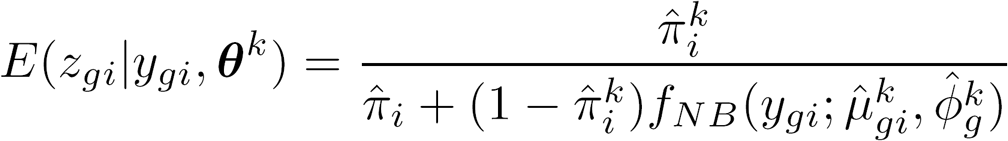
2. **M-step:** Maximize *Q*^*k*+1^ to update parameter estimates.

1. The parameters for the count component 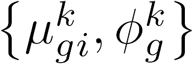 are updated using the edgeR framework^2^ by incorporating observation-level weights 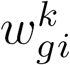 Equation (1)). In principle, any negative binomial software tool that allows for weights can be used in this step, but we have found edgeR to provide accurate and fast results. Note, we use gene-wise dispersion estimates and we do not use shrinkage within the EM-algorithm.
2. The mixture parameters 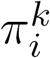 are updated with a logistic regression model of 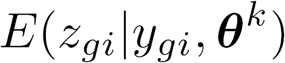 on the effective library size of a sample 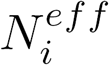 to estimate the expected probability of zero-inflation for a cell *i*

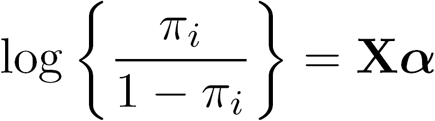

where **X** is the model matrix containing an intercept and the effective library size 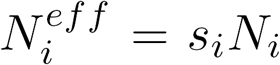 with *N_i_* the library size and *s_i_* the normalization factor for cell *i* as estimated with a global scaling normalization procedure. The normalization procedures used in this manuscript are implemented as default options in the zingeR package. However, zingeR can work with any global scaling normalization procedure when providing user-defined normalization factors as an optional argument. Optionally, other predictors can be used in the zero-excess model that are associated with zero-inflation, for example the batch effects in our case study. To gain power, the zero-excess model may also incorporate a measure for the gene's average expression which is also linked to zero abundance (Supplementary Figure 5). However, we found that this model deteriorates FDR control and consider this a topic for further research.
3. Iterate step 1 and 2 until convergence of the data log-likelihood.

### Speeding up the EM algorithm

Estimating the negative binomial parameters {*μ_gi_, ϕ_g_*} for the count component is the most computer intensive step of the EM algorithm. In order to reduce computational burden we developed an EM-algorithm that only estimates the count component parameters after iterative convergence between the mixture parameters 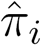 and the posterior probabilities *w_gi_*. The algorithm can be described in pseudocode as follows

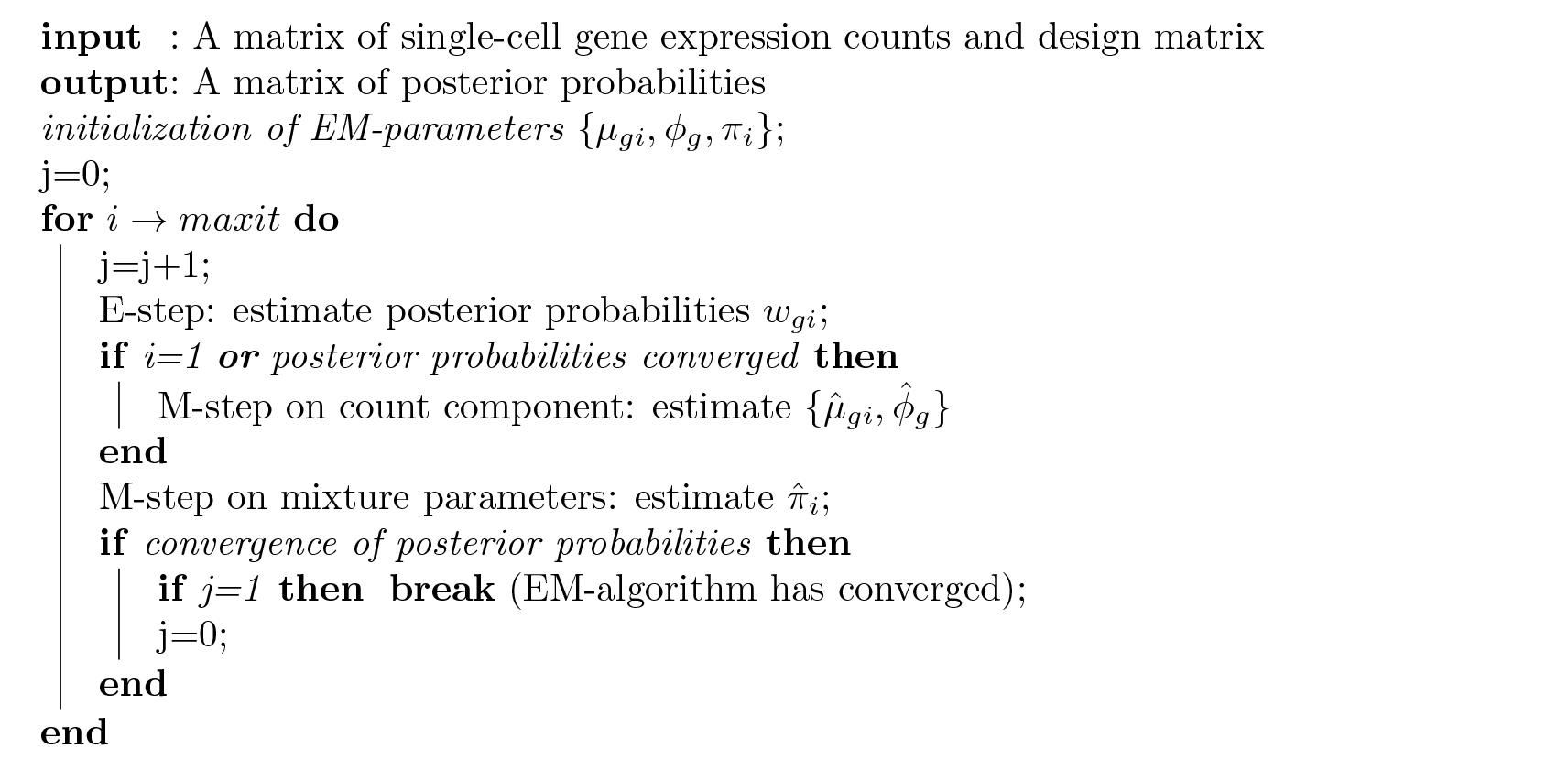

The fast version of the EM-algorithm provides very similar distributions for the posterior probabilities on the Islam and Trapnell datasets (Supplementary Figure 6).

### Inference

We only consider statistical inference on the count component of the mixture distribution. For zingeR_edgeR, we refit the models with the posterior probabilities of the converged algorithm and adopt approximate empirical Bayes shrinkage of the dispersion. Downweighting is accounted for in the statistical test by adjusting the degrees of freedom of the null distribution accordingly. More specifically, we reintroduce the moderated F-test in the edgeR package where the denominator residual degrees of freedom for a particular gene are adjusted by the extent of zero-inflation that was identified for this gene, i.e. 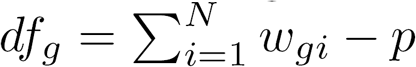 where *df_g_* are the degrees of freedom for gene *g, w_gi_* the posterior probabilities for gene *g* in sample *i* and *p* the number of model parameters estimated in the NB GLM.

We also extended the DESeq2 package to accommodate for zero-inflation by providing the option to use observation-level weights in the parameter estimation steps. DESeq2’s default normalization procedure requires geometric means of counts, which are zero for genes with at least one zero count. This limits the number of genes that can be used for normalization in scRNA-seq applications. We therefore implemented the normalization method suggested in the phyloseq package,^32^ which calculates geometric means for a gene by only using its positive counts, so genes with zero counts could still be used for normalization purposes. The phyloseq normalization procedure can now be adopted by specifying the type `poscounts` in the DESeq2 `estimateSizeFactors` function. To account for downweighting of excess zeros, we replace the Gaussian null distribution of the Wald test by a t-distribution with adjusted degrees of freedom as in the zingeR_edgeR analysis.

For limma-voom, heteroscedastic weights are estimated based on the mean-variance trend of the log-transformed counts with the voom method. The heteroscedastic weights are then multiplied with the zero-inflation weights, which are then used in a weighted linear model fit. To account for downweighting, the residual degrees of freedom of the linear model fit are adjusted before the empirical Bayes variance shrinkage and are therefore also propagated to the residual degrees of freedom from the moderated statistical tests.

For the zero-inflated methods, we use the independent filtering procedure that was developed in the genefilter package and used in DESeq2^1^ to improve performance where possible.^39^ Similar to DESeq2, we use the average expression strength (or the average fitted values) of each gene as its filter criterion and all genes with normalized mean below a filtering threshold are discarded for the multiple testing adjustment. By default, a threshold is chosen that maximizes the number of differentially expressed features.

### RNA-seq data simulation

We simulate realistic RNA-seq data based on the framework provided by Zhou et al. (2014).^28^ In brief, we estimate gene-wise means *μ_g_* and dispersions *ϕ_g_* from the Bottomly dataset^29^ and simulate RNA-seq counts according to a gene-wise negative binomial distribution where the means and dispersions are jointly sampled to respect the mean-variance relationship. We consider a two-group comparison with five biological replicates in every group. In the RNA-seq simulation, 20,000 genes are simulated according to the negative binomial distribution and fold changes are simulated according to an exponential distribution truncated at 2 as in Soneson et al. 2016.^40^ We incorporate zero-inflation by randomly replacing 5% of all counts by zeros.

### scRNA-seq data simulation

We extend the framework from Zhou et al. (2014)^28^ towards scRNA-seq applications and provide user-friendly software to simulate scRNA-seq data as part of the zingeR R package. The user can input a real scRNA-seq dataset to extract feature-level parameters for generating scRNA-seq counts. Library sizes for the simulated samples are by default resampled from the real dataset but they can also be specified. The simulation framework models positive and zero counts separately using a hurdle model. For the positive counts, expression fractions 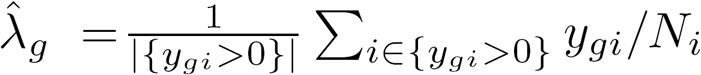 are empirically estimated and dispersions *ϕ_g_* are estimated according to a zero-truncated negative binomial (ZTNB) distribution. The zero abundance *p_gi_* of a gene *g* is modelled as a function of an interaction between its expression intensity (in terms of average log counts per million *A_g_*) and the sequencing depth of the sample *i* using a semiparametric additive logistic regression model, motivated by dataset-specific associations observed in real scRNA-seq datasets (Supplementary Figures 3, 5). The simulation paradigm jointly samples the gene-wise estimated parameters 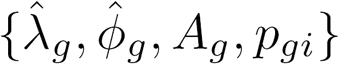 to retain gene specific characteristics present in the original dataset. We use the expected probability on zero counts 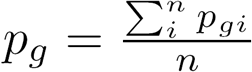 to introduce zero counts by simulating from a binomial process. Positive counts are then simulated according to a ZTNB distribution with mean 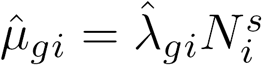 and dispersion 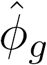 with 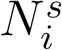 the simulated library size for sample *i*. The framework acknowledges both gene-specific characteristics as well as broad dataset-specific associations across all genes and provides realistic scRNA-seq data for method evaluation.

We evaluate performance based on both the Islam^16^ and a subset of the Trapnell^30^ dataset. The count table from the Islam dataset was downloaded from the Gene Expression Omnibus with accession number GSE29087. The Islam dataset considers 44 embryonic fibroblasts and 48 embryonic stem cells in mouse. Negative control wells are removed and 11,796 genes with at least five positive counts are retained for analysis. For the simulation, we replicate a dataset with two groups of 40 samples. The Trapnell dataset is downloaded from the preprocessed single-cell data repository conquer (http://imlspenticton.uzh.ch:3838/conquer).^24^ We only use a subset of the Trapnell dataset from the 48h and 72h timepoints of differentiating human myoblasts to generate a two-group comparison. Wells that do not contain one cell or that contain debris were removed from the dataset. We use a more stringent filtering criterion for the larger Trapnell dataset and retained 24, 576 genes with at least 10 positive counts. The simulated dataset contains two conditions with 75 samples in each condition, thereby replicating the sample size of the Trapnell dataset.

### Case study

The expression data for the case study was downloaded from Supplementary Data accompanying the original paper downloaded at http://linnarssonlab.org/drg/. Non single-cells are removed and only neuronal cells are retained for analysis, resulting in the set of 622 cells that were used for the main analyses in Usoskin et al. (2014).^11^ For differential expression analysis, the picking session was included as a covariate in all models. Contrasts were defined for the model coefficients to test for mean differential expression between one cell type and the average across all other cell types. Posterior probabilities are estimated with zingeR using 500 EM iterations or until convergence. For the mock comparison, we create two conditions with 45 cells each. In every condition 15 cells from each picking session are randomly selected over all cell types, for 30 iterations. In every iteration we test for significance of the mock variable and evaluate the false positive rate by considering the proportion of p-values ≤ 0.05.

### Method comparison

We compare zingeR to state of the art RNA-seq tools edgeR (v3.19.0),^2,41^ DESeq2 (v1.17.1)^1^ and limma-voom (v3.30.13);^3^ scRNA-seq dedicated tools scde (v2.1.2),^22^ MAST (v0.933)^19^ and NODES (v0.0.0.9010);^26^ and metagenomeSeq (v1.15.4)^27^ developed to account for zero-inflation in metagenomics applications. A ZINB model is also implemented in ShrinkBayes,^42^ but the method does not scale to the typical sample sizes observed in scRNA-seq data and has many tuning parameters, which leads us to not consider the method for comparison purposes. For all methods, all genes are considered for analysis unless the method has a default filtering step (e.g. independent filtering step for DESeq2 and zingeR based on the p-values). All samples are retained for analysis except for the NODES analysis where the default filtering step on the samples was used since it would frequently run into errors otherwise. For DESeq2, we allowed for default shrinkage of the fold changes because this notably improved its performance and we disabled the default imputation step. Also, for the zingeR_DESeq2 analysis we used the new **poscounts** normalization procedure explained above. Other settings were set to the default for all other methods. Performance is compared based on false discovery proportion-true positive rate (FDP-TPR) curves using the iCOBRA package.^43^ The p-values for all methods have been corrected with the Benjamini and Hochberg FDR method,^44^ unless specified otherwise.

### Implementation

All code to reproduced the analyses reported in the paper are available at https://github.com/statOmics/zingeRPaper. Our method and the simulation framework is available as an R package zingeR: zero-inflated negative binomial gene expression in R and development will be hosted on GitHub at https://github.com/statOmics/zingeR. The package will be submitted to R/Bioconductor (http://www.bioconductor.org).

## Acknowledgements

This research was supported in part by IAP research network “StUDyS” grant no. P7/06 of the Belgian government (Belgian Science Policy) and the Multidisciplinary Research Partnership “Bioinformatics: from nucleotides to networks” of Ghent University. KVDB is supported by a Strategic Basic Research PhD grant from the Research Foundation - Flanders (FWO) no. 1S 418 16N. CS is supported by the Forschungskredit of the University of Zurich, grant no. FK-16-107. ML is supported by NIH grant no. CA142538-07. We thank Joris Meys for his advice on building the R package.

## Author contributions

LC and KVDB conceived and designed the study. LC and KVDB implemented the method and KVDB performed the analyses. ML extended the DESeq2 package. All authors contributed to the writing of the manuscript. All authors read and approved the final manuscript.

## References

1 Michael I Love, Wolfgang Huber, and Simon Anders. Moderated estimation of fold change and dispersion for RNA-seq data with DESeq2. Genome Biology, 15(12):550, dec 2014.

2 Mark D Robinson, Davis J McCarthy, and Gordon K Smyth. edgeR: a Bioconductor package for differential expression analysis of digital gene expression data. Bioinformatics (Oxford, England), 26(1):139–40, jan 2010.

3 Charity W Law, Yunshun Chen, Wei Shi, and Gordon K Smyth. voom: Precision weights unlock linear model analysis tools for RNA-seq read counts. Genome biology, 15(2):R29, jan 2014.

4 Zhong Wang, Mark Gerstein, and Michael Snyder. RNA-Seq: a revolutionary tool for transcriptomics. Nature Reviews Genetics, 10(1):57–63, jan 2009.

5 Sara Goodwin, John D. McPherson, and W. Richard McCombie. Coming of age: ten years of next-generation sequencing technologies. Nature Reviews Genetics, 17(6):333–351, may 2016.

6 Tapio Lönnberg, Valentine Svensson, Kylie R James, Daniel Fernandez-Ruiz, Ismail Sebina, Ruddy Montandon, Megan S F Soon, Lily G Fogg, Arya Sheela Nair, Urijah Liligeto, Michael J T Stubbington, Lam-Ha Ly, Frederik Otzen Bagger, Max Zwiessele, Neil D Lawrence, Fernando Souza-Fonseca-Guimaraes, Patrick T Bunn, Christian R Engwerda, William R Heath, Oliver Billker, Oliver Stegle, Ashraful Haque, and Sarah A Teichmann. Single-cell RNA-seq and computational analysis using temporal mixture modelling resolves Th1/Tfh fate bifurcation in malaria. Science immunology, 2(9), mar 2017.

7 Florian Buettner, Kedar N Natarajan, F Paolo Casale, Valentina Proserpio, Antonio Scialdone, Fabian J Theis, Sarah A Teichmann, John C Marioni, and Oliver Stegle. Computational analysis of cell-to-cell heterogeneity in single-cell RNA-sequencing data reveals hidden subpopulations of cells. Nature Biotechnology, 33(2):155–160, jan 2015.

8 Anoop P. Patel, Itay Tirosh, John J. Trombetta, Alex K. Shalek, Shawn M. Gillespie, Hiroaki Wakimoto, Daniel P. Cahill, Brian V. Nahed, William T. Curry, Robert L. Martuza, David N. Louis, Orit Rozenblatt-Rosen, Mario L. Suvà, Aviv Regev, and Bradley E. Bernstein. Single-cell RNA-seq highlights intratumoral heterogeneity in primary glioblastoma. Science, 344(6190), 2014.

9 Aleksandra A Kolodziejczyk, Jong Kyoung Kim, Jason C H Tsang, Tomislav Ilicic, Johan Henriksson, Kedar N Natarajan, Alex C Tuck, Xuefei Gao, Marc Bühler, Pentao Liu, John C Marioni, and Sarah A Teichmann. Single Cell RNA-Sequencing of Pluripotent States Unlocks Modular Transcriptional Variation. Cell stem cell, 17(4):471–85, oct 2015.

10 Li Li, Ji Dong, Liying Yan, Jun Yong, Xixi Liu, Yuqiong Hu, Xiaoying Fan, Xinglong Wu, Hongshan Guo, Xiaoye Wang, Xiaohui Zhu, Rong Li, Jie Yan, Yuan Wei, Yangyu Zhao, Wei Wang, Yixin Ren, Peng Yuan, Zhiqiang Yan, Boqiang Hu, Fan Guo, Lu Wen, Fuchou Tang, and Jie Qiao. Single-Cell RNA-Seq Analysis Maps Development of Human Germline Cells and Gonadal Niche Interactions. Cell Stem Cell, 20(6):858–873.e4, jun 2017.

11 Dmitry Usoskin, Alessandro Furlan, Saiful Islam, Hind Abdo, Peter Lönnerberg, Daohua Lou, Jens Hjerling-Leffler, Jesper Haeggström, Olga Kharchenko, Peter V Kharchenko, Sten Linnarsson, and Patrik Ernfors. Unbiased classification of sensory neuron types by large-scale single-cell RNA sequencing. Nature Neuroscience, 18(1):145–153, nov 2014.

12 Aleksandra A. Kolodziejczyk, Jong Kyoung Kim, Valentine Svensson, John C. Marioni, and Sarah A. Teichmann. The Technology and Biology of Single-Cell RNA Sequencing. Molecular Cell, 58(4):610–620, may 2015.

13 T. Nakamura, Y. Yabuta, I. Okamoto, S. Aramaki, S. Yokobayashi, K. Kurimoto, K. Sekiguchi, M. Nakagawa, T. Yamamoto, and M. Saitou. SC3-seq: a method for highly parallel and quantitative measurement of single-cell gene expression. Nucleic Acids Research, 43(9):e60–e60, may 2015.

14 Angela R Wu, Norma F Neff, Tomer Kalisky, Piero Dalerba, Barbara Treutlein, Michael E Rothenberg, Francis M Mburu, Gary L Mantalas, Sopheak Sim, Michael F Clarke, and Stephen R Quake. Quantitative assessment of single-cell RNA-sequencing methods. Nature Methods, 11(1):41–46, oct 2013.

15 Saiful Islam, Amit Zeisel, Simon Joost, Gioele La Manno, Pawel Zajac, Maria Kasper, Peter Lönnerberg, and Sten, Linnarsson. Quantitative single-cell RNA-seq with unique molecular identifiers. Nature Methods, 11(2):163–166, dec 2013.

16 Saiful Islam, Una Kjällquist, Annalena Moliner, Pawel Zajac, Jian-Bing Fan, Peter Lönnerberg, and Sten Linnarsson. Characterization of the single-cell transcriptional landscape by highly multiplex RNA-seq. Genome research, 21(7):1160–7, jul 2011.

17 Simone Picelli, Omid R Faridani, ?sa K Bj?rklund, G?sta Winberg, Sven Sagasser, and Rickard Sandberg. Full-length RNA-seq from single cells using Smart-seq2. Nature Protocols, 9(1):171–181, jan 2014.

18 Tamar Hashimshony, Naftalie Senderovich, Gal Avital, Agnes Klochendler, Yaron de Leeuw,Leon Anavy, Dave Gennert, Shuqiang Li, Kenneth J Livak, Orit Rozenblatt-Rosen, Yuval Dor, Aviv Regev, and Itai Yanai. CEL-Seq2: sensitive highly-multiplexed single-cell RNA-Seq. Genome biology, 17:77, apr 2016.

19 Greg Finak, Andrew McDavid, Masanao Yajima, Jingyuan Deng, Vivian Gersuk, Alex K. Shalek, Chloe K. Slichter, Hannah W. Miller, M. Juliana McElrath, Martin Prlic, Peter S. Linsley, and Raphael Gottardo. MAST: a flexible statistical framework for assessing transcriptional changes and characterizing heterogeneity in single-cell RNA sequencing data. Genome Biology, 16(1):278, dec 2015.

20 Arjun Raj, Charles S Peskin, Daniel Tranchina, Diana Y Vargas, and Sanjay Tyagi. Stochastic mRNA Synthesis in Mammalian Cells. PLoS Biology, 4(10):e309, sep 2006.

21 Aaron T.L. Lun, Davis J. McCarthy, and John C. Marioni. A step-by-step workflow for low-level analysis of single-cell RNA-seq data with Bioconductor. F1000Research, 5:2122, oct 2016.

22 Peter V Kharchenko, Lev Silberstein, and David T Scadden. Bayesian approach to single-cell differential expression analysis. Nature methods, 11(7):740–2, jul 2014.

23 Maria K. Jaakkola, Fatemeh Seyednasrollah, Arfa Mehmood, and Laura L. Elo. Comparison of methods to detect differentially expressed genes between single-cell populations. Briefings in Bioinformatics, page bbw057, jul 2016.

24 Charlotte Soneson and Mark D. Robinson. Bias, Robustness And Scalability In Differential Expression Analysis Of Single-Cell RNA-Seq Data. bioRxiv, 2017.

25 Davis J McCarthy, Yunshun Chen, and Gordon K Smyth. Differential expression analysis of multifactor RNA-Seq experiments with respect to biological variation. Nucleic acids research, 40(10):4288–97, may 2012.

26 Debarka Sengupta, Nirmala Arul Rayan, Michelle Lim, Bing Lim, and Shyam Prabhakar. Fast, scalable and accurate differential expression analysis for single cells. bioRxiv, 2016.

27 Joseph N Paulson, O Colin Stine, Héctor Corrada Bravo, and Mihai Pop. Differential abundance analysis for microbial marker-gene surveys. Nature methods, 10(12):1200–2, sep 2013.

28 Xiaobei Zhou, Helen Lindsay, and Mark D Robinson. Robustly detecting differential expression in RNA sequencing data using observation weights. Nucleic acids research, 42(11):e91, jun 2014.

29 Daniel Bottomly, Nicole A R Walter, Jessica Ezzell Hunter, Priscila Darakjian, Sunita Kawane, Kari J Buck, Robert P Searles, Michael Mooney, Shannon K McWeeney, and Robert Hitzemann. Evaluating gene expression in C57BL/6J and DBA/2J mouse striatum using RNA-Seq and microarrays. PloS one, 6(3):e17820, jan 2011.

30 Cole Trapnell, David G Hendrickson, Martin Sauvageau, Loyal Goff, John L Rinn, and Lior Pachter. Differential analysis of gene regulation at transcript resolution with RNA-seq. Nature biotechnology, 31(1):46–53, jan 2013.

31 Stephanie C Hicks, Mingxiang Teng, and Rafael A Irizarry. On the widespread and critical impact of systematic bias and batch effects in single-cell RNA-Seq data. bioRxiv, 2015.

32 Paul J. McMurdie and Susan Holmes. phyloseq: An R Package for Reproducible Interactive Analysis and Graphics of Microbiome Census Data. PLoS ONE, 8(4):e61217, apr 2013.

33 Aaron T L Lun, Karsten Bach, and John C Marioni. Pooling across cells to normalize single-cell RNA sequencing data with many zero counts. Genome biology, 17:75, apr 2016.

34 Dominic Grün, Lennart Kester, and Alexander van Oudenaarden. Validation of noise models for single-cell transcriptomics. Nature Methods, 11(6):637–640, apr 2014.

35 Simon Anders and Wolfgang Huber. Differential expression analysis for sequence count data. Genome Biology, 11(10):R106, 2010.

36 GK Smyth. Linear models and empirical bayes methods for assessing differential expression in microarray experiments. Statistical applications in genetics and molecular biology, 3(1): 1–26, 2004.

37 Bradley Efron. Large-Scale Inference. Empirical Bayes Methods for Estimation, Testing and Prediction. Cambridge University Press, New York, 2010.

38 Mark D Robinson and Alicia Oshlack. A scaling normalization method for differential expression analysis of RNA-seq data. Genome Biology, 11(3):R25, 2010.

39 Richard Bourgon, Robert Gentleman, and Wolfgang Huber. Independent filtering increases detection power for high-throughput experiments. Proceedings of the National Academy of Sciences of the United States of America, 107(21) :9546–51, may 2010.

40 Charlotte Soneson, Michael I. Love, and Mark D. Robinson. Differential analyses for RNA-seq: transcript-level estimates improve gene-level inferences. F1000Research, 4:1521, feb 2016.

41 Davis J McCarthy, Yunshun Chen, and Gordon K Smyth. Differential expression analysis of multifactor RNA-Seq experiments with respect to biological variation. Nucleic acids research, 40(10):4288–97, may 2012.

42 Mark A van de Wiel, Maarten Neerincx, Tineke E Buffart, Daoud Sie, and Henk MW Verheul. ShrinkBayes: a versatile R-package for analysis of count-based sequencing data in complex study designs. BMC Bioinformatics, 15(1):116, 2014.

43 Charlotte Soneson and Mark D Robinson. iCOBRA: open, reproducible, standardized and live method benchmarking. Nature Methods, 13(4):283–283, mar 2016.

44 Yaov Benjamini and Yosef Hochberg. Controlling the False Discovery Rate: A Practical and Powerful Approach to Multiple Testing. Journal of the Royal Statistical Society. Series B (Methodological), 57(1):289–300, 1995.

